# Adaptive optics allows 3D STED-FCS measurements in the cytoplasm of living cells

**DOI:** 10.1101/584870

**Authors:** Aurélien Barbotin, Silvia Galiani, Iztok Urbančič, Christian Eggeling, Martin Booth

**Affiliations:** Department of Engineering Science, University of Oxford, Parks Road, Oxford OX1 3PJ, UK; MRC Human Immunology Unit, MRC Weatherall Institute of Molecular Medicine, University of Oxford, Oxford OX3 9DS, UK; Wolfson Imaging Centre Oxford, MRC Weatherall Institute of Molecular Medicine, University of Oxford, Oxford OX3 9DS; Institute of Applied Optics, Friedrich-Schiller-University Jena, Max-Wien Platz 4, 07743 Jena, Germany; Leibniz Institute of Photonic Technology e.V., Albert-Einstein-Strasse 9, 07745 Jena, Germany; “Jožef Stefan” Institute, Jamova 39, SLO-1000 Ljubljana (Slovenia)

## Abstract

Fluorescence correlation spectroscopy in combination with super-resolution stimulated emission depletion microscopy (STED-FCS) is a powerful tool to investigate molecular diffusion with sub-diffraction resolution. It has been of particular use for investigations of two dimensional systems like cell membranes, but has so far seen very limited applications to studies of three-dimensional diffusion. One reason for this is the extreme sensitivity of the axial (3D) STED depletion pattern to optical aberrations. We present here an adaptive optics-based correction method that compensates for these aberrations and allows STED-FCS measurements in the cytoplasm of living cells.

## 1. Introduction

Fluorescence correlation spectroscopy (FCS) provides measurements of molecular diffusion based on the fluctuations of a fluorescence signal induced by molecules moving in and out of an observation volume (1, 2). From the autocorrelation function (ACF) of these fluctuations, it is possible to extract the average transit time of a fluorophore and the average number of molecules in the observation volume. A classical implementation of FCS makes use of confocal optics, but more recently FCS was demonstrated in subdiffraction volumes using super-resolution stimulated emission depletion (STED) microscopy (STED-FCS) (3). STED-FCS is based on confocal microscopy, where (as in usual STED microscopy implementations) an additional (STED) laser with a central intensity zero is added to the diffraction-limited confocal fluorescence excitation laser to inhibit fluorescence emission everywhere but at the very centre. Depending on the power of the STED laser, the size of the effective fluorescence observation spot can thus be tuned to sub-diffraction scales. Depending on the choice of depletion pattern, STED can either increase lateral resolution (2D STED, using a ring-shaped intensity pattern) or mainly axial resolution (3D STED, using a so-called "bottle beam") (4, 5). 2D STED-FCS has proven to be a tool of value to investigate two-dimensional diffusion in systems like the cellular membrane (4, 5), but faces severe limitations in the case of 3D diffusion. 2D STED-FCS in 3D environments constrains the lateral resolution and assumes uniform diffusion properties along the optical axis. In this configuration, the estimation of both average number of molecules in the observation volume and transit times is complicated by varying axial cross-section of the beam and biased by out-of-focus contributions (5, 6), requiring either specific fitting models (6, 7), background subtraction using separation of photons by lifetime tuning (SPLIT) (8), or stimulated emission double depletion (STEDD) (9).

In principle, the 3D depletion pattern should greatly simplify the analysis and interpretation of STED-FCS measurements of 3D diffusion. However, the main factor limiting the use of 3D STED-FCS is the extreme sensitivity to optical aberrations (10–12) that can fill up the central intensity minimum or distort the focus, reducing brightness and resolution (13) and ultimately leading to an amount of noise precluding reliable STED-FCS measurements.

These problems can be overcome using methods of adaptive optics (AO), in which adaptive elements, such as spatial light modulators (SLMs) correct aberrations introduced by the specimen (14). AO has already been proven successful to increase both resolution and signal to noise ratio (SNR) in STED microscopy (15–17) and FCS (18–21), but not in STED-FCS. We present here an implementation of AO for STED-FCS that allowed us to correct the aberrations induced by a refractive index mismatch and by misalignments caused by mechanical drift in both freely diffusing dyes in solution and in cells, providing accurate and sensitive 3D STED-FCS measurements in the cytoplasm of living cells.

## 2. Materials and methods

### A. Microscope layout

We used a custom STED microscope built around a RESOLFT microscope from Abberior Instruments as described in previous publications (22), and as sketched in figure 1. In short, the excitation focus was created by focusing a 640 nm pulsed laser with a 100×/1.4 oil immersion objective (Olympus UPLSAPO). Excitation power was set to 8.6 μW in solution and to 6 μW in cells. We employed a 755 nm pulsed laser (Spectra-Physics Mai Tai, pulse-stretched by a 40-cm glass rod and a 100-m single-mode fibre) with a repetition frequency of 80 MHz as fluorescence depletion or STED laser, modulated in phase using a spatial light modulator (SLM) (Hamamatsu LCOS X10468-02, figure 1). The STED laser power was set to 16 mW for the aberration correction procedures, and varied between 6 and 60 mW for STED-FCS measurements. Fluorescence light was collected by the objective, filtered by a pinhole and measured with an avalanche photodiode (APD). The microscope was controlled by the software Imspector (Abberior Instruments). The SLM was controlled by a bespoke python software and was used for both phase mask generation and aberration correction. AO feedback loops were implemented in python using the Imspector python interface.

**Fig. 1.**
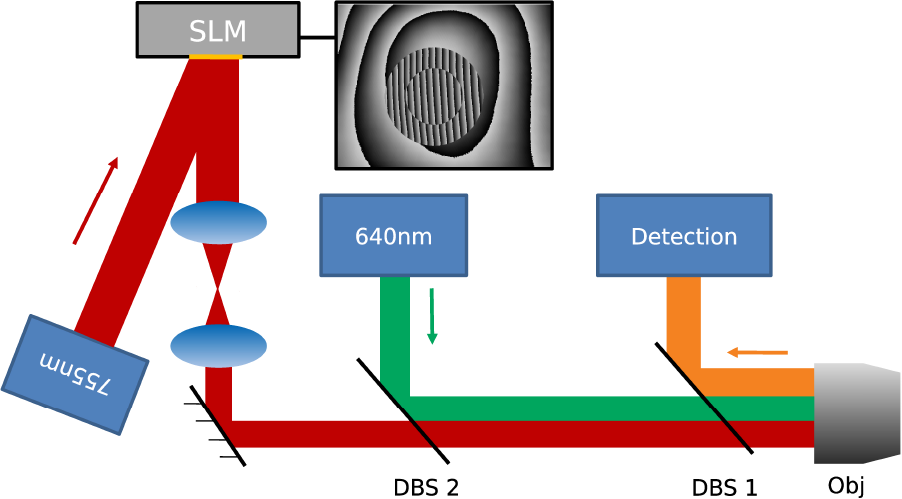
Schematic of STED-FCS setup with 755 nm STED and 640 nm excitation lasers (blue boxes and red and green beam paths), fluorescence detection (blue box and orange beam path), lenses (blue ellipses), mirrors (black lines), spatial light modulator (SLM), dichroic beam splitters (DBS 1 and 2), objective lens (Obj). The inset shows a phase mask applied on the spatial light modulator (SLM), created from a flatness correction, a blazed grating to transport the pupil in an off-axis hologram, a central *π* phase shift to create a 3D STED depletion pattern and a phase aberration correction.

System aberrations in the depletion path were removed by imaging a specimen of scattering gold beads using the sensorless method. Image quality was assessed using image standard deviation. Coalignment between excitation and depletion was ensured by imaging a specimen of 40 nm crimson beads (Abberior), following the method described in (23), using instead a wavelet-based image quality metric (24).

Data acquisition was performed using the software Imspector. Fluorescence intensity time traces were recorded with sampling frequency of 1 MHz.

### B. FCS

Auto correlation functions (ACFs) *G*(*τ*) were obtained by correlating fluorescence intensity time traces *I*(*t*) according to equation 1 using the python package *multipletau* (25).

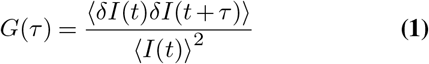

where ⟨.⟩ is the operator averaging over time (*t*) and *δI*(*t*) = *I*(*t*)−⟨*I*(*t*)⟩.

From ACFs, the average number of molecules in the observation volume and their diffusion coefficient can be estimated. A general form of the ACF model can be written as:

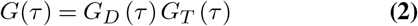

with *G*_*D*_ (*τ*) and *G*_*T*_ (*τ*) being the ACF contributions arising from intensity fluctuations due to diffusion of molecules and electron transitions to the triplet state, respectively.

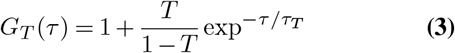

where *T* is the average triplet amplitude and *τ*_*T*_ is the triplet correlation time. Assuming a 3D Gaussian intensity profile of the effective observation volume (which is applicable even for 3D STED-FCS (3, 5)), the diffusive part of the ACF can be expressed as:

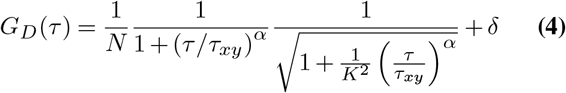

where *N* is the apparent average number of molecules in the observation volume, *δ* is an offset, *τ*_*xy*_ is the lateral transit time, *α* is a factor that characterises deviations from the Brownian diffusion model, and *K* is the aspect ratio of the Gaussian-assumed effective observation volume (or effective optical point-spread function (PSF)) defined as *K* = *ω*_*z*_/*ω*_*xy*_, where *ω*_*z*_ is the axial and *ω*_*xy*_ the lateral 1/e^2^ radius.

### C. Data fitting

A considerable challenge of 3D STED-FCS is data analysis. An ideal model would need to consider independent shrinking of lateral and axial dimensions of the effective observation volume with the STED power, but realistic signal-to-noise ratio of acquired STED-FCS curves, especially from within cells, typically preclude reliable independent fitting of the two transit times (3, 5). To this end, certain simplifications of the model are required to gain robustness in analysis. In previous implementations of 3D STED-FCS, the FCS curves were fitted using a 3D Gaussian model, assuming a constant lateral transit time, measured independently in confocal mode (3, 5). However, this does neglect that also in the 3D STED modality the lateral diameter of the PSF is decreased by up to 30 % (see figure 2, a), leading to an in principle 2-fold reduction in lateral transit time.

We therefore developed a different fitting method that takes into account also this lateral shrinking of the PSF, which was empirically determined from images of immobilised 40 nm crimson beads. Fitting the axial and lateral intensity profiles with a Gaussian function allowed determination of the variations of lateral and axial diameters of the PSF (given as full width at half-maximum (FWHM) of the profiles) at different STED laser powers (figure 2, a). This empirical relationship between the lateral resolution improvement and the aspect ratio *K*, which we fitted with an exponential function (figure 2, b), permits description of the PSF (and thus effective observation volume) with one parameter instead of two. The diffusive parts of the FCS curves were therefore fitted according to equation 4, where the aspect ratio *K* varied with the ratio of the lateral transit times *τ*_*xy,STED*_/*τ*_*xy,confocal*_ (from STED and confocal recordings) according to our empirical calibration.

We verified the performance of the new fitting approach and compared it to the previous method for fitting 3D STED-FCS data of dyes in solution and in cytoplasm. The general size of the observation volumes as determined by our novel fitting approach were similar to those determined by fitting the aspect ratio *K* only, making our results direcly comparable to those from earlier literature. Yet, as highlighted before, the novel approach well accounted for the present slight lateral reduction of the observation volume with increasing STED power (Fig. 2). We also found our novel method to return more realistic values when fitting non-ideal curves with small contributions of spurious correlating signal, often appearing during measurements in cells. (see figure 1 of supplement 1). Purely confocal data were fitted by fixing the aspect ratio *K* to a reasonable value of 4, as in a previous implementation of 3D STED-FCS (3), and optimising *τ*_*xy*_. We set a higher confocal aspect ratio for STED-FCS than determined from imaging to account for aberrations affecting the excitation beam. Triplet correlation times were determined independently and fixed to 12 μs in solution and 5 μs in cells.

**Fig. 2.**
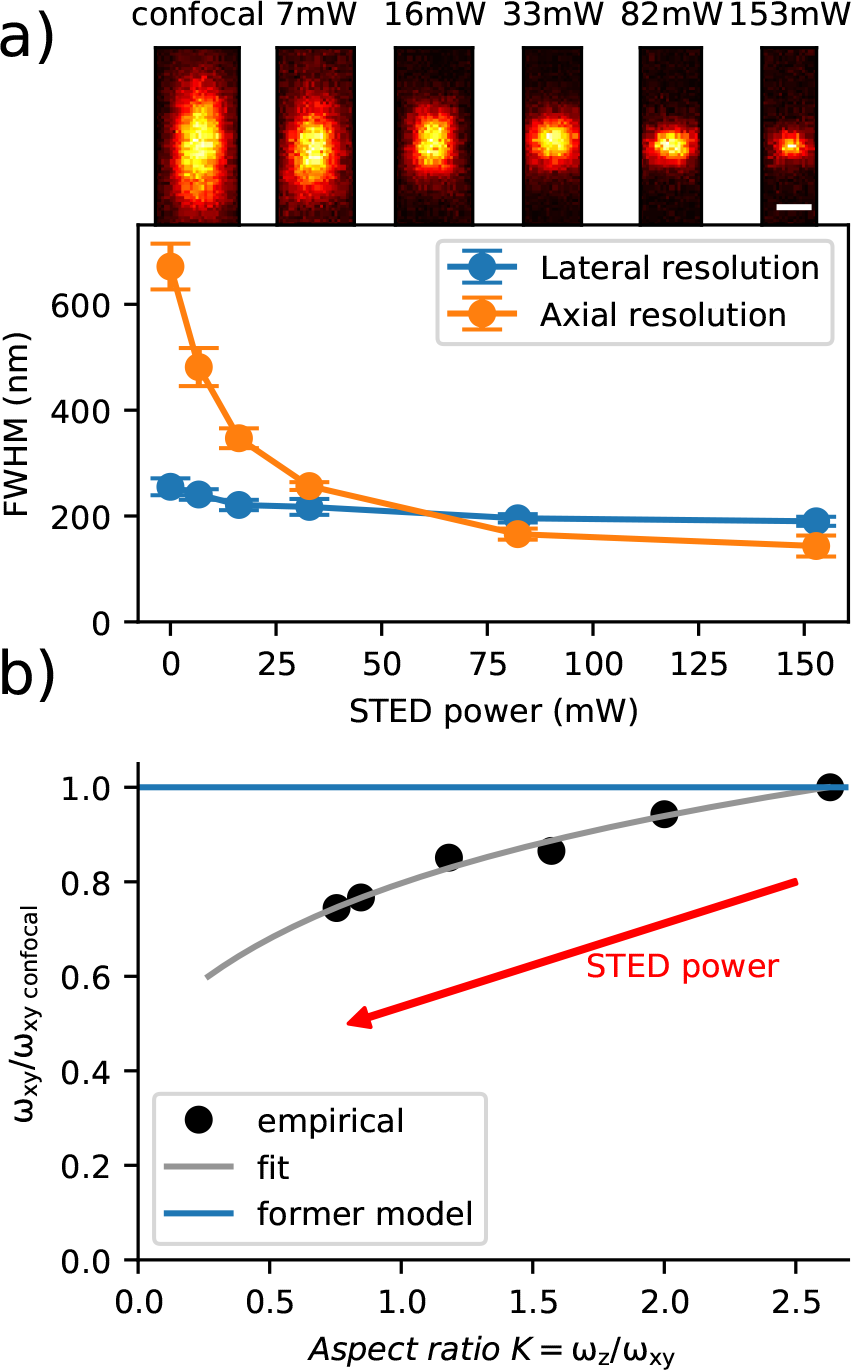
Variations in the shape of the STED laser focus with STED laser power. **a)**: Determination of lateral and axial resolutions at different STED laser powers using images of fluorescent beads. Top: images of fluorescent beads acquired at different STED laser powers (scale bar 200 nm). Bottom: Gaussian fits to intensity profiles over images of individual beads returning axial and lateral resolution variations with STED laser power (mean +/− s.d., n = 10 beads). **b)**: Values of the lateral beam waist *ω*_*xy*_ (normalized to the value *ω*_*xy,confocal*_) as a function of the aspect ratio *K* = *ω*_*z*_/*ω*_*xy*_ as determined from the bead images of **a** (black dots, empirical) were fitted with an exponential function (grey curve) describing the variations of the shape of the observation volume with STED laser power to design a novel STED-FCS fitting model. The previous model neglecting the reduction in lateral dimension (blue line) is also plotted for comparison.

### D. Samples. Dye samples

A solution of freely diffusing dye molecules was prepared by diluting Abberior Star Red NHS dyes (Abberior) in a 1:1 water:glycerol solution to a concentration of 50 nM. Glycerol was used to increase the viscosity of the medium and decrease the diffusion speed of the dyes, which otherwise diffuse too fast for reliable assessment of average number of molecules in the observation volume and transit times with STED-FCS.

#### Cells

Human fibroblasts (GM5756T, Moser, Baltimore, USA) were maintained in a culture medium consisting of DMEM with 4500 mg glucose/L, 110 mg sodium pyruvate/L supplemented with 10% fetal calf serum, glutamine (2 mM) and penicillin-streptomycin (1%). The cells were cultured at 37 °C/ 5% CO_2_. Cells were grown in a 35 mm imaging dish with an ibidi polymer coverslip bottom (ibidi GmbH, Germany), and transfected with a plasmid expressing a fusion protein of GFP and SNAP-tag using Lipofectamine 3000 transfection reagent (Invitrogene, Carlsbad, USA). 24 hours after transfection, the cells were incubated together with SNAP-Cell 647-SiR (New England Biolabs (UK) Ltd., Hitchin, UK) and washed twice in culture medium after 40 min incubation, with a waiting time between washings of 20 min. Finally the culture medium was substituted with L-15 medium (Sigma-Aldrich, Dorset, UK) and each sample was visualized at 37°C for no longer than 2 hours.

## 3. Adaptive optics

We used sensorless AO (26) to remove aberrations affecting the depletion beam only, as described in figure 3a.

### A. Aberration modes

Sensorless AO requires wavefront decomposition into a set of modes that are individually corrected. We chose the Zernike polynomials as modes, as is the case in most sensorless AO microscopes (18, 27). In the case of aberrations affecting STED depletion beams, it was demonstrated that certain modes induce not only a deformation of the depletion pattern, but also a shift of the central intensity minimum (12, 15, 28). We measured experimentally the shifts induced by each mode and removed them with tip, tilt and defocus as described in (15). Instead of Zernike defocus, which induces axial shifts but also deformations, we used the exact expression of defocus in high numerical aperture (NA) systems (29):

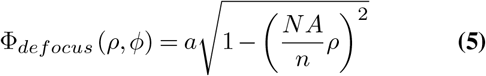

where Φ_*def ocus*_ is the circular phase function in the back focal plane (BFP) of the objective inducing defocus, *ρ* and *ϕ* are the cylindrical coordinates of the BFP, *NA* the numerical aperture of the objective, *n* the refractive index of the immersion medium, and *a* is an amplitude factor. High NA defocus was not normalised like other Zernike modes; however throughout this paper we will refer to the amount of high NA defocus introduced as radians rms as a means of convenience. We set the value of the amplitude factor *a* to ensure that the shift induced by a given amount of defocus is comparable to that induced by tip or tilt, and that consequently a given amount of defocus and tip or tilt affect STED-FCS parameters in a comparable way. The shifts of the depletion pattern induced by tip, tilt and high NA defocus can be found in table 1.

**Fig. 3.**
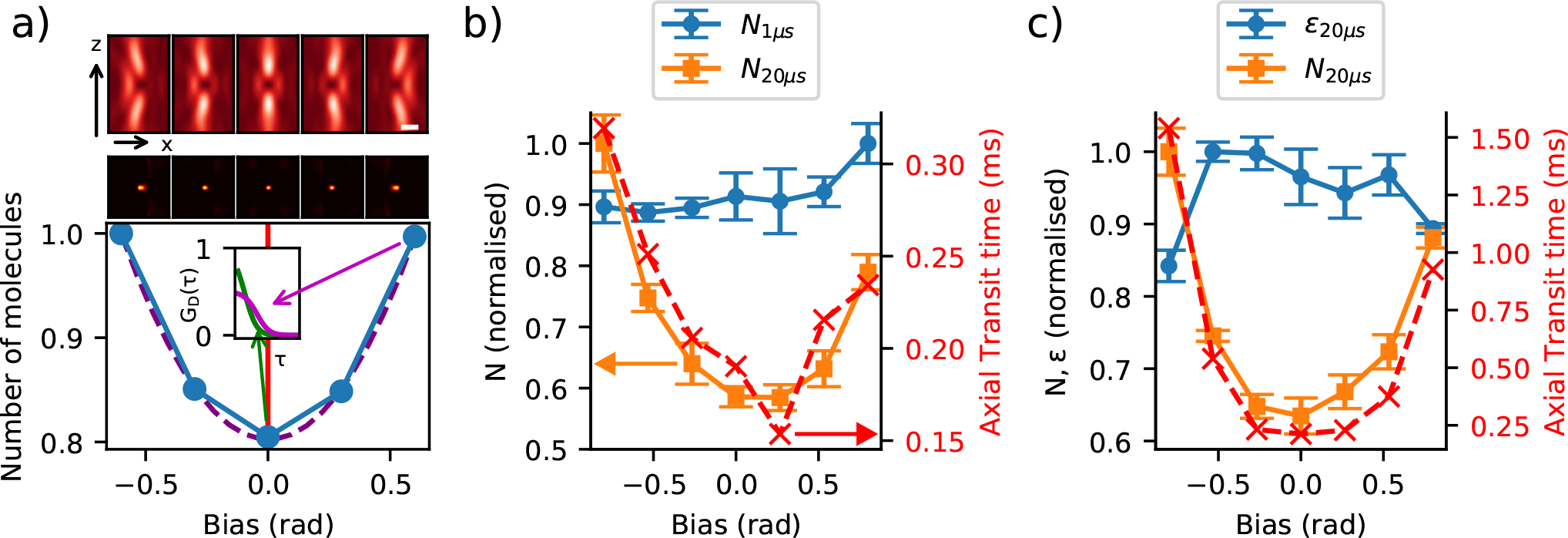
Finding an experimental metric for AO 3D-STED-FCS. **a**: Principle of our experimental procedure. A set of aberration modes (bias) are applied to the STED laser beam using the SLM creating “bottle beam” foci of different quality (top: respective simulated focal STED laser intensity pattern, middle: simulated effective observation volume. Scalebar: 400 nm). Bottom: Sketch of expected outcome; metric values (blue) against introduced bias and quadratic fit (dashed purple line) for determining the optimum (vertical red line), and (inset) sketch of expected FCS curves *G*_*D*_(*τ*) for optimal (green) and maximum biased (purple) conditions highlighting the expected aberration-introduced decrease in amplitude and increase in transit time. **b**,**c**: Experimental data from AO 3D-STED-FCS: calculated values of number of molecules (*N*) and molecular brightness (*ϵ*) for different binning times as labeled (orange, blue points, left y-axis) as well as axial transit time *τ*_*z*_ (red cross, right y-axis) (mean +/− s.d., n=5 points) against introduced bias (tilt (b) and horizontal coma (c) in rad). STED laser power = 16 mW.

**Table 1.** Theoretical spatial shifts induced by 1 radian rms of tip, tilt and high NA defocus

| Mode | Axis | Shift (nm/rad) |
| --- | --- | --- |
| Tip | x | 172 |
| Tilt | y | 172 |
| High NA defocus | z | 339 |

It is critical in STED-based microscopy to reduce the number of measurements by as much as possible to limit photo-bleaching. Because low-order Zernike modes are the most commonly encountered in biological specimens (30), we decided to correct only the first four Zernike modes (astigmatism and coma) as well as primary and secondary spherical aberrations.

Our STED microscope included a single adaptive element in the STED laser beam path only, and therefore did not correct aberrations affecting the excitation beam and the detection path. In certain cases, when aberration correction is performed solely in the STED laser beam, misalignments can occur between the excitation and STED laser beams, deteriorating the quality of the effective observation volume (17). A solution to that is to use a deformable mirror to remove aberrations affecting all beam paths, at the cost of an additional experimental complexity. Instead, we treated tip, tilt and defocus as aberration modes and optimised them along with the other modes to provide in-sample realignment. This approach also allows to correct for thermal and mechanical drift that can affect the coalignment of the excitation and depletion beams.

### B. Metric design

The choice of a suitable quality metric is essential to a successful sensorless AO implementation. In confocal FCS, the overall signal intensity (average photon count) was successfully employed (18). However, in the case of STED microscopy, different aberration modes affect the depletion pattern adversely, either increasing or decreasing the overall signal intensity (11, 12, 15, 28). The optimal metrics for STED-FCS should report on the size and/or quality of the effective observation volume. This information could be obtained from parameters (*τ*_*xy*_, *τ*_*z*_, *N*) extracted from fitting the FCS curves, but the computational cost for this would make the optimisation procedure impractically slow. Besides, the fitting procedure relies on the assumption that the shape of the observation volume is Gaussian, which is not verified during the aberration correction procedure.

We instead explored higher moments of the signal intensity (or photon counts) distributions, which can be used to estimate the average number of molecules in the observation volume (*N*) and the fluorescence count rate per molecule (i.e. molecular brightness, *ϵ*) (31) following the equations:

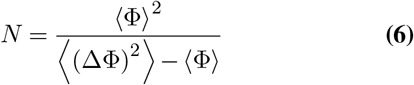

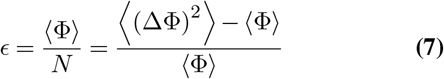

where Φ(*t*) is the detected photon count rate at time *t*, and ⟨(ΔΦ)^2^⟩ is the variance of the photon count distribution.

A low degree of aberration in the STED laser provides an optimal effective observation spot with highest possible molecular brightness *ϵ* and lowest average number of molecules *N*. Therefore, these moment values provide an accurate measure for aberration correction and as such are suited to be metrics for AO. Molecular brightness was successfully used as a metric for AO in conventional confocal FCS (20) and was demonstrated to outperform the overall photon count rate in certain situations (21).

To select the best quality metric for AO 3D-STED-FCS, we determined values of the *ϵ* and *N* for freely diffusing Abberior Star Red dyes in a 1:1 water:glycerol solution while inducing various amounts of aberration (or bias) affecting the quality of the 3D-STED laser focus (“bottle beam”). We recorded 10 s photon count time traces for different amounts of aberrations introduced by the SLM (see figure 3, a). Each 10 s photon count trace was split into five sub-traces each of 2 s duration and values of *ϵ* and *N* calculated using equations 6 and 7. Their outputs were compared to the variations in transit times from fitting the corresponding ACFs, which served as a reference.

Photon count time traces were originally recorded with a time binning of 1 μs. At this low integration times, intensity fluctuations are not only caused by molecular diffusion, but also influenced by Poisson noise and by molecular blinking due to transient transitions into the dark triplet states(32). It is known that further binning of the photon count time traces alleviates these effects(at the expense changing absolute values of *ϵ* and *N* (33)), and hence (since we did not want to introduce challenging corrections for triplet state dynamics and Poisson noise) further binning of the time traces greatly improved the aberration measurement accuracy (figure 3, b). The choice of binning factor was empirical and corresponded to a tradeoff between the necessity to remove the fast fluctuations described above and the necessity to keep fluctuations induced by molecular motion large enough.

Comparing both metrics to the axial transit time *τ*_*z*_, we found that the molecular brightness *ϵ* is not a good metric for 3D-STED-FCS, since it measures two competing effects: molecular brightness can decrease both if the central intensity minimum of the depletion beam fills in (lower quality) or shrinks (smaller observation volume, better quality). On the contrary, assuming a constant molecular concentration, the average number of molecules *N* in the observation volume provides a direct measure of the size of the observation volume and hence exactly corresponds to the parameter that needs to be optimised (figure 3 c).

## 4. Results

### A. AO improves the performance of STED-FCS in solution

We first set up and evaluated our aberration correction routine for 3D-STED-FCS on a solution of freely diffusing dyes at a depth of 3 μm (typical depth for FCS measurements of molecules diffusing in cytosol) To highlight the effect of aberration correction, we used an oil immersion objective with an intentional refractive index mismatch to the aqueous sample. For each aberration mode described in section A, we acquired intensity timetraces with an acquisition time of 3 s for 7 different bias aberrations introduced by the SLM (see figure 3, a). Each timetrace was re-binned at 20 μs before calculating the average number of molecules *N* in the observation volume from the detected photon count rate as the quality metric, as described in section 3.B. The resulting quality metric curve (*N* versus introduced bias) was fitted with a quadratic function to determine its minimum (smallest *N* corresponds to the smallest observation volume), subsequently defined as the optimal correction. We refer to this whole procedure as a round of aberration correction (see figure 3, a).

**Fig. 4.**
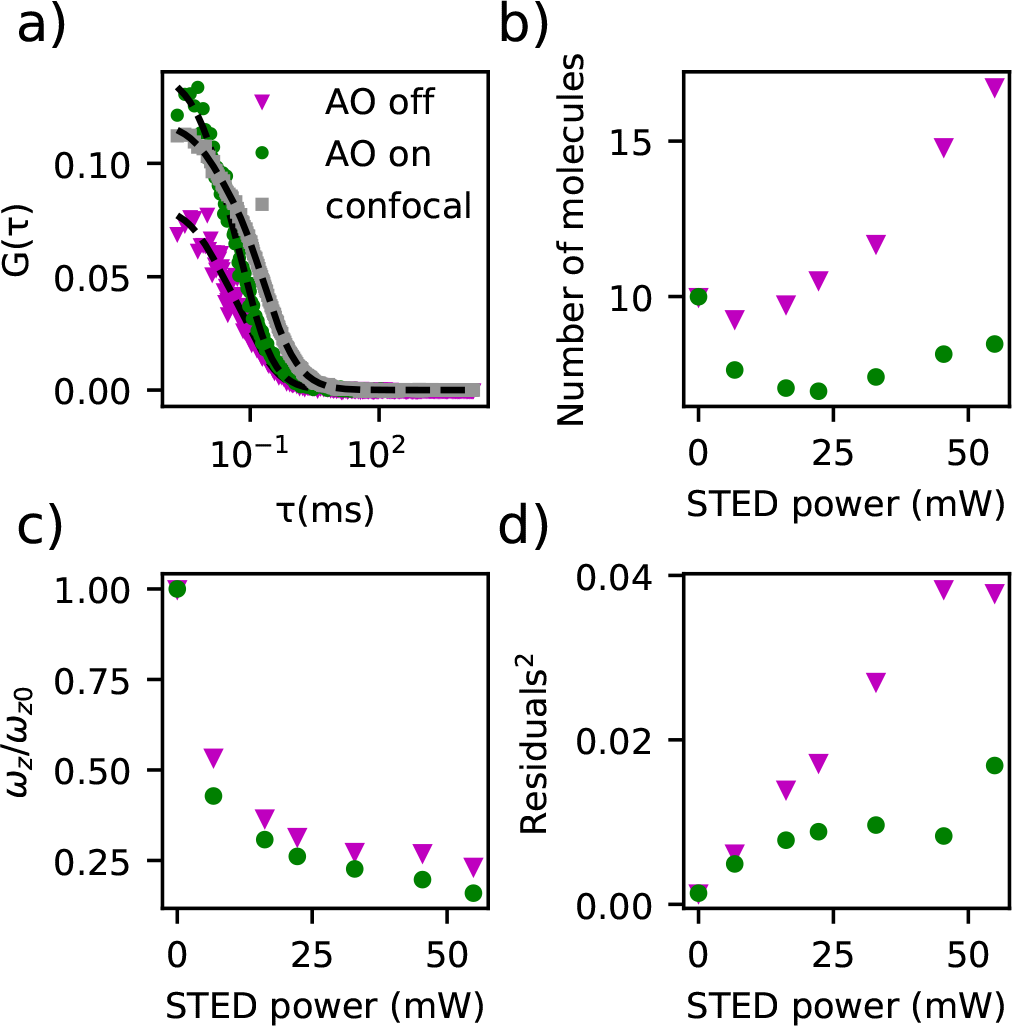
Aberration correction for 3D-STED-FCS diffusion measurements of Abberior Star Red in water:glycerol solution, measured 3 μm above the coverslip. **a**: FCS curves (*G*(*τ*)) for confocal mode only (grey squares) and with a STED laser power of 55 mW without (magenta triangles, AO off) and with (green dots, AO on) AO correction, and with fits (equation 2, dashed black lines). **b**-**d**: Resulting values of **b)** average number of molecules *N* in the observation volume, **c)** ratio *ω*_*z*_/*ω*_*z*0_ of axial diameters of the observation volume (*ω*_*z*_, equation 4) for the respective STED laser power (*ω*_*z*_) and for the confocal recordings (*ω*_*z*0_ at 0mW STED laser power), and **d)** the squared sum of residuals from the fit to the data (normalised by the square of the ACF amplitude, as a measure of the noise and deviation from the fit model) as a function of the STED laser power without (magenta triangles) and with (green dots) AO correction.

We found that a single round of aberration correction was sufficient to correct for coma, astigmatism, tip, tilt and defocus, and that more rounds of aberration correction were necessary to sufficiently remove primary and secondary spherical aberrations. This is consistent with previously reported AO applications in STED microscopy imaging (15).

The maximal amount of aberration introduced in a correction round represents a tradeoff between measurement precision (higher with low aberration amplitude) and sensitivity to noise (lower with high aberration amplitude). We achieved robust convergence of the optimisation using a maximum aberration amplitude of 0.8 radian for correction of tip, tilt, astigmatism and coma as well as for the first three rounds of spherical and defocus correction. We then performed two more rounds of spherical aberration correction with an amplitude of 0.6 radians to ensure an optimal correction.

After having determined the optimal set of aberration correction parameters, we evaluated the impact of AO on 3D-STED-FCS measurements by acquiring a series of 30s-long intensity time-traces at various STED powers, with and with-out AO correction (figure 4). FCS curves were calculated and fitted as described in section B to determine values for the observation volume parameters (lateral *ω*_*xy*_ and axial *ω*_*z*_ diameters, equation 4) and the average number of molecules *N* within the observation volume. Aberration correction induced a dramatic increase in the amplitude of the FCS curves (figure 4, a), which implies a reduction in the determined values of *N* (figure 4, b), as would be expected for a less aberrated observation volume. AO also improved the resolution, as highlighted by a reduced axial extent of the observation volume (figure 4, c). Residuals from fitting FCS curves (normalized to an amplitude of 1) were used as a measure of both SNR (larger SNR result in less noisy curves and thus lower residuals)) and conformity to the fitting model. We found AO to significantly decrease the fitting residuals (figure 4, d).

**Fig. 5.**
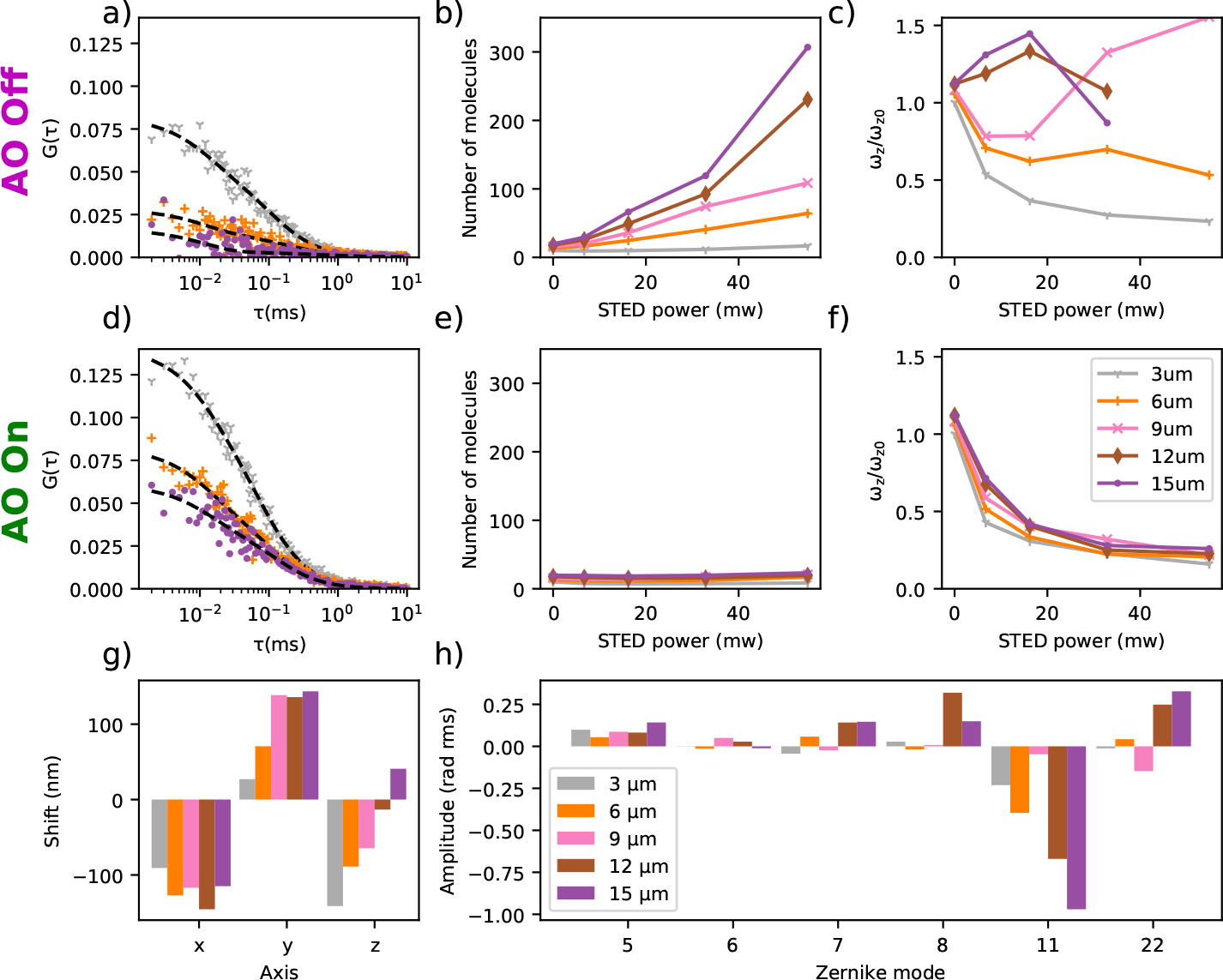
AO correction of depth-induced aberrations in 3D STED-FCS measurements: diffusion of Abberior Star Red in a water:glycerol solution. **a,d** Representative FCS data *G*(*τ*) at a STED laser power of 55 mW (dashed lines: fits to the data), **b,e** number of molecules *N* as determined from FCS data recorded for different STED laser powers, and **c,f** ratio *ω*_*z*_/*ω*_*z*0_ of axial diameters of the observation volume (equation 4) as determined from the FCS data at different STED laser powers (*ω*_*z*_) and confocal recordings (*ω*_*z*0_) with (**d-f**) and without (**a-c**) AO aberration correction and for different depths as detailed in the colour legend in panel **f**. **g)** Depth-dependent spatial repositioning of the depletion pattern using tip, tilt and defocus (x,y,z). **h)** Determined depth-dependent correction values for aberrations modes. Zernike modes were numbered following the convention defined by Noll [30].

In previous implementations of 3D STED-FCS, the determined values of the average number of molecules *N* in the observation volume was found to increase monotonically with STED laser power (3), a result apparently inconsistent with the expected decrease in observation volume induced by the STED laser. This increase in determined values of *N* was due to increasing contributions of undepleted non-correlated out-of-focus background (4, 5) that damp the amplitude of the FCS curves and (since from theory the amplitude is inversely proportional to *N*) lead to an increase of determined values of *N* (34). We found here that the improved SNR introduced by AO allows a considerable reduction in the relative contribution of uncorrelated background fluorescence and consequently a reduction in the resulting values of *N* (figure 4, b).

### B. AO extends the range of observation depths for STED-FCS in solution

We then increased the observation depth to test the capability of our system to correct depth-induced aberrations. We increased the depth positioning of the laser focus and thus of the observation volume in steps of 3 μm, from 6 to 15 μm. At each depth, we performed one round of aberration correction for astigmatism, coma, tip and tilt and two rounds of correction for defocus, primary and secondary spherical. We found it necessary to increase the acquisition times for aberration measurement at increasing depths to compensate for the loss of signal caused by aberrations affecting the excitation and detection paths. We increased acquisition times from 3 s at 6 μm to 12 s at 15 μm. The AO procedure at each depth was performed from the correction determined at the previous depth, i.e. we started the aberration correction procedure at 12 μm with the aberrations determined at 9 μm.

After having determined the optimal correction at each depth, we recorded and correlated 30 s intensity time traces, at four different STED powers with and without AO, and in confocal mode (figure 5, a and d). Fitting the ACFs, we estimated apparent average number of molecules in the observation volume (figure 5, b and e) and axial resolution improvement (figure 5, c and f). As expected, the apparent average number of molecules in the observation volume increased much more without aberration correction due to an increase in uncorrelated background and reduction in signal levels. We also found that at depths larger than 6 μm, estimation of observation volumes without AO was compromised (the low SNR of ACFs precluded reliable fitting of transit times), while AO showed a consistent decrease in the size of the observation volume with STED laser power even at large depth (figure 5).

**Fig. 6.**
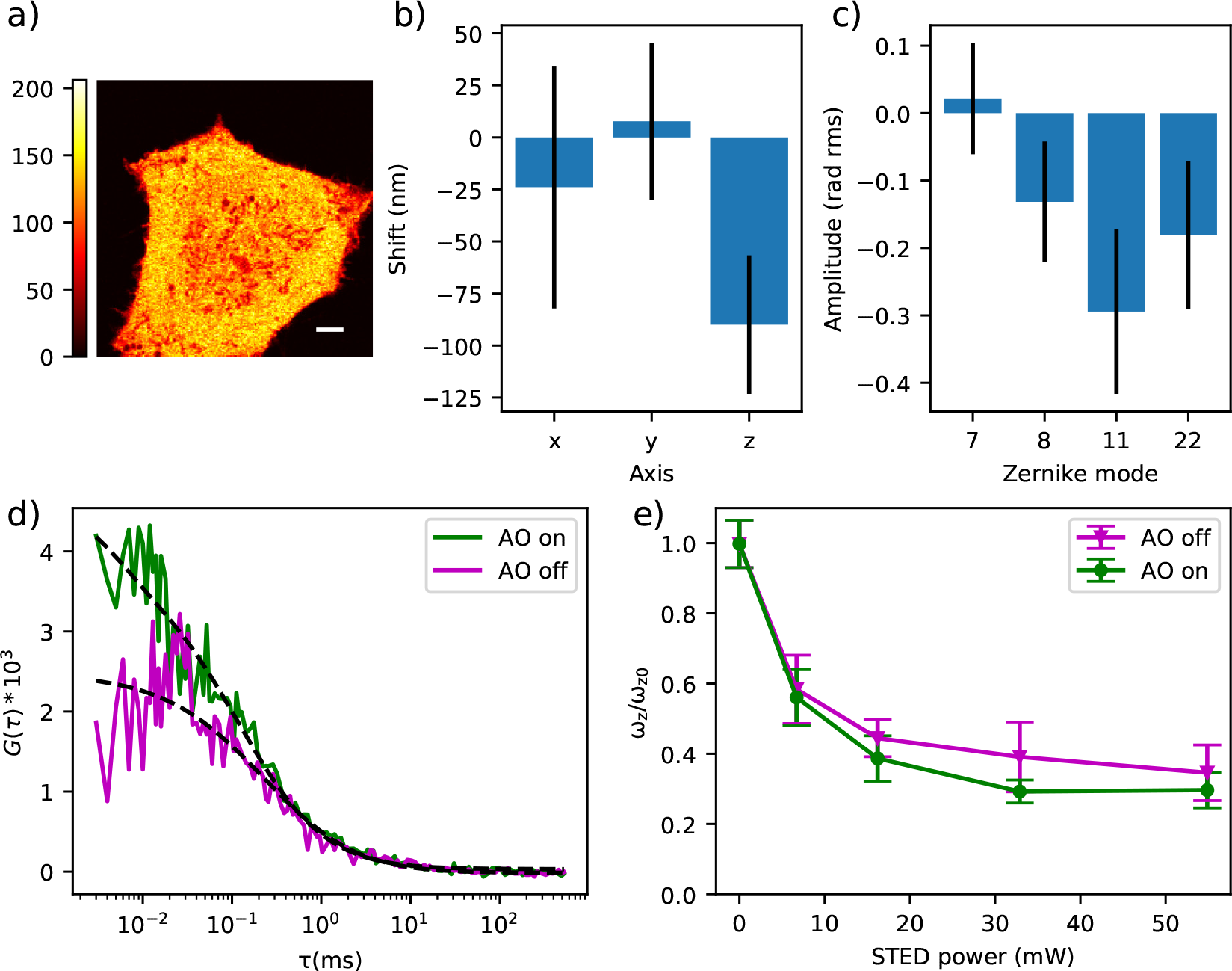
Aberration correction of live-cell 3D STED-FCS measurements: cytoplasmic 647-SiR diffusion. **a**: Confocal image of a representative cell where aberrations were corrected for (scalebar: 5 μm). **b**: Spatial shifts of the depletion pattern and **c** aberration values measured in cells (mean +/− s.d, n=17 measurements in 15 cells, 1-2 measurements per cell). **d**: Representative FCS curves with and without aberration correction at a STED power of 33 mW (dashed lines: fits to the data). **e**: Ratio *w*_*z*_/*w*_*z*0_ of axial diameters of the observation volume (equation 4) as determined from the FCS data at different STED laser powers (*w*_*z*_) and confocal recordings (*w*_*z*0_) (mean +/− s.d., from 16-19 curves per datapoint).

The retrieved aberration corrections indicate that the major source of the depth-induced deterioration of the observation volume was spherical aberration caused by the refractive index mismatch between the immersion oil and the solution (Zernike modes 11 and 22 in figure 5, h). The measured values of primary and secondary spherical aberrations varied linearly with penetration depth. At 9 μm penetration depth, a rogue measurement skewed the estimation of spherical modes, which we attempted to correct by doing two more rounds of spherical aberration correction. As a result, the amount of spherical aberrations determined at this depth did not follow the general trend (figure 5, h).

At every depth, repositioning of the depletion beam using tip, tilt and defocus was necessary to compensate for a mixture of different effects (figure 5, g), such as imprecision of the instrument calibration procedure, thermal and mechanical drift, and spatial shifts of the excitation and depletion beams induced by aberrations. We verified that the lateral shifts could not be the consequence of the coma (Zernike modes 7 and 8) we had to correct for, so they most probably originated from general mechanical drifts in the microscope.

### C. Aberration correction in living cells

We finally demonstrated aberration correction in the cytoplasm of living cells, performing STED-FCS measurements of freely diffusing dyes in human fibroblasts (figure 6, a).

Aberration correction was performed using seven data points per mode with a 4 s acquisition time each, and a maximum bias of 0.8 radians rms. To reduce photobleaching and total light exposure, as well as to increase experimental efficiency, it is beneficial to minimise the number of these AO correction measurements. For this reason we reconsidered the need to correct all the aberration modes we had applied in solutions. Spherical aberrations and coma are introduced by a refractive index mismatch and a tilted sample, respectively, and are therefore expected to be present in microscopy experiments with cells. So can be astigmatism (35), but its values that we measured in a set of preliminary STED-FCS experiments were negligible (less than 0.2 radians rms). Besides, astigmatism has little effect on the 3D STED depletion beam (12). For these reasons we decided not to correct astigmatism in STED-FCS measurements with cells.

With this approach we performed a series of AO STED-FCS experiments with a cytoplasmic dye in cells at depths between approximately 2 and 6 μm. The structure of aberrations (figure 6, b and c) expectedly confirmed the spherical aberrations to be the dominant modes.

Based on these results we also decided not to correct for the secondary spherical aberration, as considerable cross-talk between the primary and secondary spherical modes required at least two rounds of corrections at a minor additional improvement in the signal quality. With this final scheme, we performed aberration correction in the cytosol of a cell at a depth of approximately 4 μm. Once the correction was determined, we acquired a series of FCS curves in the vicinity of the area where aberration correction was performed, at four different STED powers and in confocal mode. At each STED laser power, 40 FCS curves were acquired with an acquisition time of 5 s, 20 with the AO correction and 20 without. FCS curves were acquired alternatively with AO on and off to ensure that any difference observed between the two modalities was not due to photobleaching or time.

Effects of photobleaching within each acquisition were mitigated by applying post-processing correction, using the local averaging method (36), and curves that did not converge towards 0 at longer lag times were considered as affected by artefacts and were manually discarded.

Due to the crowded environment of the cell cytoplasm, molecules can undergo anomalous subdiffusion. We determined by fitting confocal ACFs that diffusion in the cytosol of our cells was well described by a parameter *α* = 0.75 (equation 4), in accordance with values previously reported in the cytoplasm of cells (37).

Results obtained are presented in figure 6. As in solution, we found that relatively low amounts of aberrations were responsible for a sensible SNR reduction in STED-FCS measurements without AO, reducing the ACF amplitude (figure 6, d), and decreasing spatial resolution (figure 6, e).

## 5. Discussion and conclusions

We proved here that AO can bring significant improvements to STED-FCS experiments in 3D by means of correction of residual system aberrations, correction of the effects of thermal and mechanical drift that could shift the depletion pattern by more than 100 nm (see figure 5, g), and most of all correction of aberrations induced by a refractive index mismatch. The latter could be reduced, although not entirely removed, by using a water immersion objective, but this approach would come at the price of a lower NA and so of a lower resolution and signal levels. Besides, the refractive index of cells can significantly deviate from that of water (38, 39) and vary from cell to cell, which can be much more efficiently handled by the adaptive aberration correction method than manual re-adjustments of the correction ring on the objective.

The substantial improvement in SNR as well as in resolution brought by adaptive optics allowed STED-FCS measurements with a high spatial resolution, allowing an up to 10-fold reduction in observation volume compared to confocal recordings (see figure 1 of Supplement 1). This is the best reduction in observation volume size obtained to our knowledge with 3D STED-FCS in an aqueous solution. Smaller observation volumes could only be obtained in organic solvents minimising the refractive index mismatch (see table 2). Adaptive optics also allowed a significant increase of the maximum focussing depth when using an oil immersion objective, from 6 μm without AO to more than 15 μm with AO. Finally, it allowed significant improvements in signal levels and resolution in the cytoplasm of cells, which paves the way to a wealth of new applications.

**Table 2.** Reduction in observation volume size obtained from different STED-FCS approaches for 3D diffusion.

| Medium | Method | Reduction in observation volume size |
| --- | --- | --- |
| Aqueous | SPLIT-FLCS (8) | 4 |
|  | 3D STED-FCS (3) | 5 |
|  | STEDD (9) | 5 |
|  | 2D STED-FCS (5) | 10 |
|  | AO 3D STED-FCS (See figure 1 of supplement 1) | 10 |
| Organic | 3D STED-FCS (5) | 15 |
|  | 2D STED-FCS (5) | 25 |

## Acknowledgments

The authors thank Katharina Reglinski for providing us with plasmids and Falk Schneider for his useful advice.

