## Supplement 1 for "Adaptive optics allows 3D STED-FCS measurements in the cytoplasm of living cells"

### Supplement 1: Comparison between fitting focal volume and fitting aspect ratio

We compared our new fitting method, using pre-calibrated relationship between the focal volume and shape, with the previously established method , which used a fixed (confocal) transit time and fitted the aspect ratio only (1). We compared the focal volumes estimated from each fitter on a series of FCS curves acquired in a solution of freely diffusing Abberior Star Red dyes in a water:glycerol solution at a depth of 3 $\mu$m (figure 1, a). We found that the focal volumes estimated with each method were simillar. Besides, we did not find a significant difference between the fiting quality in each case, as each fitting method produced simillar residuals (figure 1, b). Differences between the two methods appeared when fitting data acquired in living cells, where the heterogeneous environment can induce consequent variations in the shapes of ACFs. Since the aspect ratio has little effect on the final shape of the model, fitting aspect ratio only can easily diverge, leading to absurd values, even for curves with only slight spurious contributions (figure 1, c and d). Fitting the entire focal volume is much more stable and allows to fit a wider variety of curves.

1. Lars Kastrup, Hans Blom, Christian Eggeling, and Stefan W. Hell. Fluorescence fluctuation spectroscopy in subdiffraction focal volumes. *Physical Review Letters*, 94(17):1–4, 2005. ISSN 00319007. doi: 10.1103/PhysRevLett.94.178104.

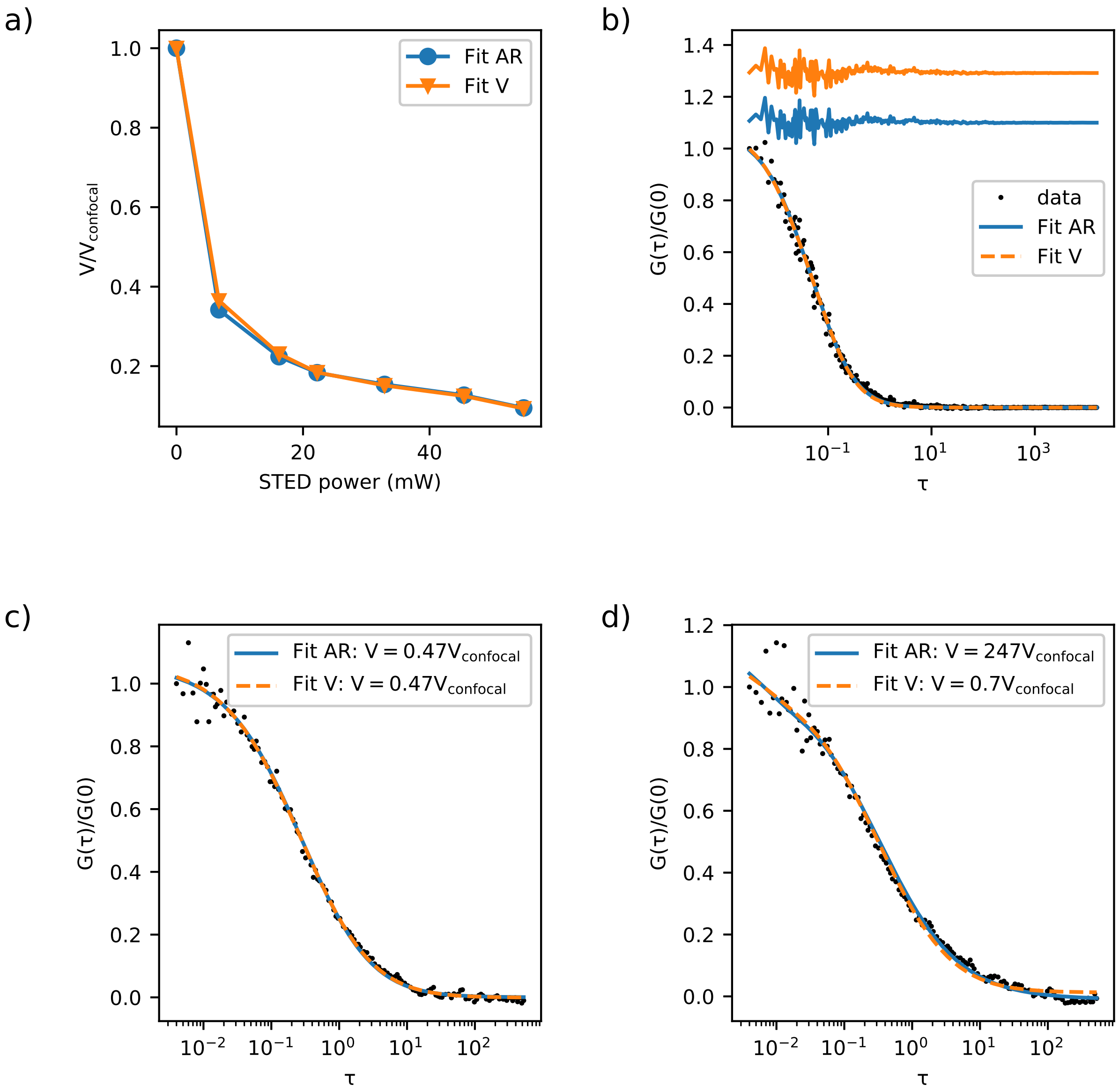


**Fig. 1.** Comparison between fitting methods for estimation of STED-FCS parameters in solution (**a-b**) and in cells (**c-d**). **a**: estimation of focal volumes of STED-FCS experiments in solution with adaptive correction, by either fitting the aspect ratio only (blue, circles) or fitting the entire volume (orange, triangles). **b**: Comparison of both fitters on a STED-FCS curve obtained in solution at a STED power of 55 mW. Residuals are plotted above the FCS curve. **c** and **d**: Fitting curves acquired in cells at a STED power of 7 mW with both models. **c**: exemplary curve that can be fitted with both fitters. Focal volumes were both equal to $0.47V_{confocal}$. **d**: exemplary curve leading to a fitting artefact when fitting aspect ratio only and not when fitting the volume. Focal volumes determined by each method were respectively equal to $247V_{confocal}$ and $0.7V_{confocal}$.
